# Single-cell transcriptomics identifies a p21-activated kinase important for survival of the zoonotic parasite *Fasciola hepatica*

**DOI:** 10.1101/2024.03.26.586785

**Authors:** Oliver Puckelwaldt, Svenja Gramberg, Sagar Ajmera, Janine Koepke, Christos Samakovlis, Simone Haeberlein

**Affiliations:** Institute of Parasitology, Justus Liebig University Giessen, Giessen, Germany; Cardio-Pulmonary Institute (CPI), Giessen, Germany

**Keywords:** Single-cell transcriptomics, scRNAseq, flatworms, liver flukes, *Fasciola hepatica*, drug target, p21-activated kinase, TRPM, stem cells

## Abstract

Knowledge on the cell types and cell-specific gene expression of multicellular pathogens facilitates drug discovery and allows gaining a deeper understanding of pathogen biology. By utilizing single-cell RNA sequencing (scRNA-seq), we analyzed 19,581 cells of a globally prevalent parasitic flatworm, the liver fluke *Fasciola hepatica*, which causes a neglected tropical disease and zoonosis known as fascioliasis. We identified 15 distinct clusters, including cells of the reproductive tract and gastrodermis, and report the identification and transcriptional characterization of potential differentiation lineages of stem cells within this parasite. Furthermore, a previously unrecognized ELF5- and TRPM_PZQ_-expressing muscle cell type was discovered, characterized by high expression of protein kinases. Among these, the p21-activated kinase PAK4 was essential for parasite survival. These data provide novel insight into the cellular composition of a complex multicellular parasite and demonstrate how gene expression at single-cell resolution can serve as a resource for the identification of new drug targets.

## Introduction

Infections with parasitic helminths pose a global health challenge. As many of these diseases affect humans and animals alike, they are of major importance considering “One Health” initiatives^1^. Fascioliasis is caused by liver flukes such as *Fasciola hepatica*, a parasitic flatworm heavily affecting livestock industry. Here, it causes a huge economic burden by reducing growth and milk yield^2^. Along with other food borne diseases, fascioliasis is recognized by the World Health Organization as a neglected tropical disease (NTD). It is estimated that up to 17 million people are infected worldwide^3^ and 90 million are at risk of infection^4^. The increasing number of reports on parasites being resistant to the commonly used drug triclabendazole^5,6^ and a lack of treatment alternatives or effective vaccines^7^ motivates basic research on liver flukes aiming at the identification of novel drug targets.

*F. hepatica* has a complex life cycle, which includes an intermediate snail host and a mammalian as final host. The adult worms reside within the bile duct of the final host, where they shed tens of thousands of eggs per day in order to reproduce^8^. The eggs are released into the environment during defecation, followed by hatching of miracidia that infect snails. Within the snail, the parasite multiplies by asexual reproduction. Infected snails eventually shed cercariae, which encyst upon aquatic vegetation, where they can remain infectious for months. After oral ingestion of infectious metacercariae by the final mammalian host, newly excysted juveniles hatch from the cyst and migrate to the liver parenchyma, where the immature worms feed and grow into adults. Adult worms can persist for decades, hinting at an outstanding longevity of these worms^9,10^.

Insights into the biology of *F. hepatica* have been significantly advanced by the implementation of various omics technologies^11^. By using bulk RNA-seq, it was possible to identify genes being transcribed in different life stages^12^, to uncover the response to anthelmintics^13,14^, investigate interactions with the immune system^15^, and assess the role of the nervous system in development^16^. While being an important tool to investigate the dynamics of transcriptional gene expression, bulk RNA-seq naturally does not allow conclusions at single-cell resolution. Bulk analysis may also complicate the detection of rare genes or dynamics in rare cell types due to overrepresented cell types or highly expressed genes. The advent of single-cell RNA sequencing (scRNA-seq) opened a new opportunity to investigate the molecular biology of multicellular organisms^17^. By profiling hundreds to thousands of cells in one experiment, this technology allows the identification and characterization of cell types and their characteristic genes even on a whole-organism level, if single-cell suspensions are accessible. This untargeted method promises unprecedented insights into the molecular biology especially of non-model organisms such as parasites, for which many other methodologies are still unavailable. In parasitic and free-living relatives of *F. hepatica,* specifically schistosomes and planarians, scRNA-seq technologies recently boosted the identification of novel cell-type markers^18,19^ and the characterization of transcription factors in developmental trajectories^20^. By focusing on cell types with vital function for a parasite, scRNA-seq data may also facilitate the identification of new drug targets.

Protein kinases (PKs) have gained attention as promising drug targets in parasites such as cestodes, filaria and other trematodes^21^. For *F. hepatica*, we recently provided evidence of several druggable PKs^22^. PKs regulate most of the core biological processes, like signal transduction, cell cycle or motility^23^. Considerable effort was put into the development of inhibitors of human PKs for use in several diseases, leading to 52 already approved drugs^24^ and several more in clinical trials^25^. Drug repurposing has been discussed as a valuable approach for the identification of new treatment options against NTDs, as there are fewer risks involved regarding efficacy and safety considerations^26^. The high number of available PK inhibitors and evidence for the druggability of kinases in parasitic worms make PKs attractive targets, potentially also for treating fascioliasis. A question to be answered is: which type of PKs can we consider important for pathogen survival, e.g. based on their expression in particular cell types?

In this study, we profiled the gene expression of adult *F. hepatica* at single-cell resolution using the 10X Chromium workflow. In order to achieve this, we established a cell dissociation protocol coupled with fluorescence activated cell sorting (FACS) to obtain a viable cell suspension. We uncovered several different cell populations, their characteristic marker genes, and identified several PKs with enriched expression in distinct cell types. Targeting one PK using a small-molecule inhibitor validated its suitability as drug target. This work presents the first single-cell atlas for this family of parasites and will serve as a resource for future biomedical research as well as basic understanding of pathogens.

## Results & Discussion

### Determination of nuclei number of *F. hepatica* adults shows cell density differences along anterior-posterior axis

Basic metrics on the number of cells in multicellular organisms are helpful in planning scRNA-seq experiments, but such metrics are unknown for liver flukes. In order to determine the total cell count and to assess the distribution of cells throughout the parasite, we quantified the nuclei within sections of adult worms. To exclude a potential bias depending on the tissue area, we utilized frontal sections (Fig 1 A) as well as transversal sections from representative areas of the worm (anterior part 1 and 2 and posterior part) (Fig 1 B). The total number of nuclei per worm was extrapolated from the nuclei counts per section. Independent of the section plane, we arrived at a total nuclei number of around 17 million within the parasite (Fig 1 C). It is to be noted that this nuclei number is only an approximation for the total cell number, as the multi-nucleated nature of the syncytial surface of the parasite, the tegument, or the fused rosette during spermatogensis^27^ do not allow a one-to-one translation.

**Fig 1.**
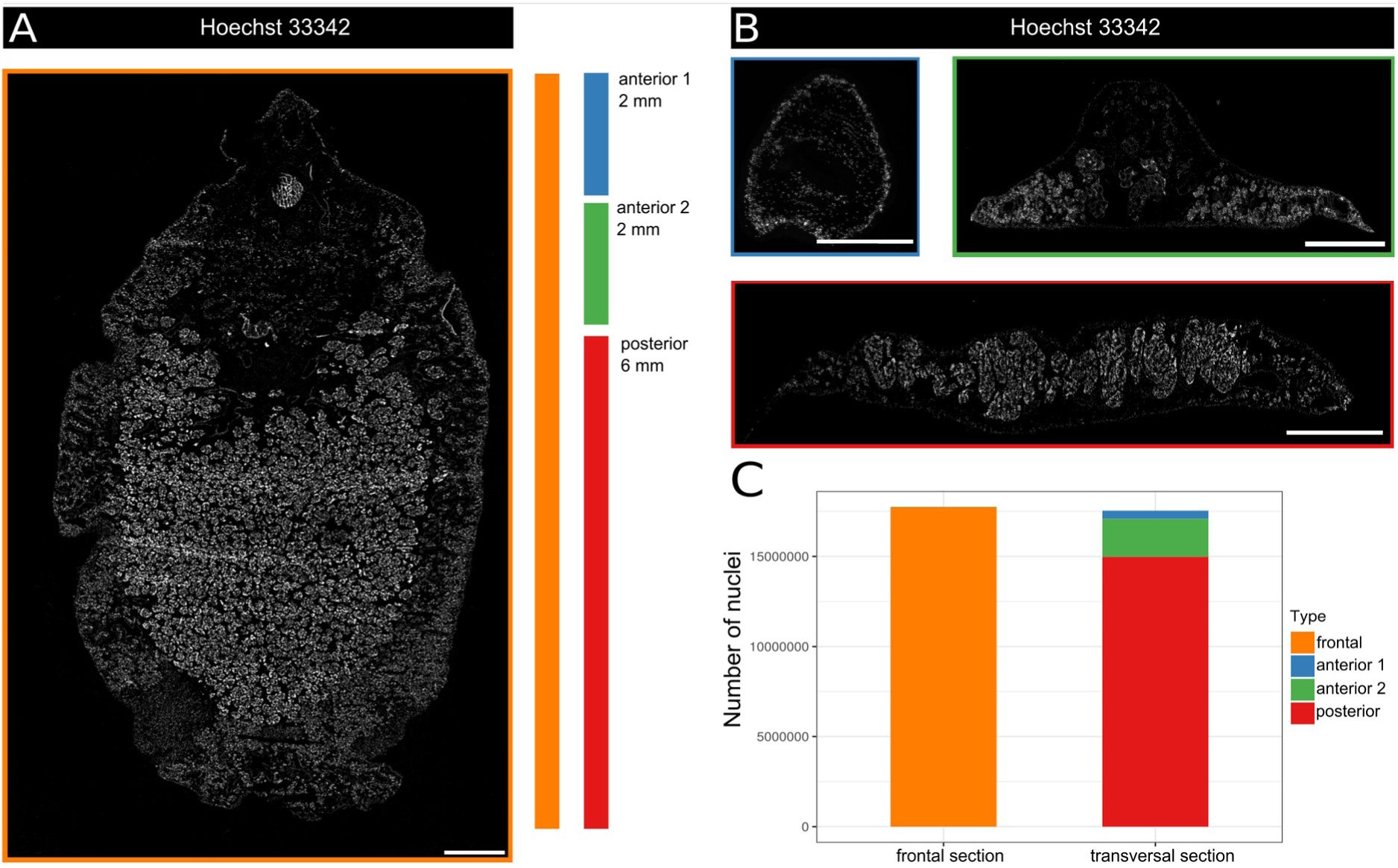
*Fasciola hepatica* is structured in regions with different cellular densities. **A** Representative frontal section of an adult worm with nuclei stained using Hoechst 33342. Color bars mark the worm regions selected for transversal sectioning. Scale bar: 1000 µm. **B** Representative images of transversal sections with nuclei stained with Hoechst 33342. Top from left to right: sections from the anterior 1 and anterior 2 regions. Bottom: section from the posterior region. Scale bar: 1000 µm. **C** Stacked bar plot indicating the extrapolated total number of nuclei derived from either frontal or transversal sections.

Compared to the two anterior parts used for quantification, most of the cells were located within the large posterior part of the worm, which contains the male reproductive tissue as well as the vitellarium, creating a higher total cell count. This not only arises from the sheer size of these tissues, in addition, the male reproductive organ as well as the vitellarium also have a high cellular density (Fig 1 A). This disproportional tissue and cell distribution over the body axis bears the risk that cells of the overabundant reproductive organs might overrun some of the rarer cell types, which are hence not captured in a subsequent 10X workflow. Based on the cellular composition, we therefore decided to cut the worm into two parts for subsequent scRNA-seq experiments: a posterior part and an anterior part, which is enriched for proportionately underrepresented cell types.

### scRNA-seq captures cells of major tissue types of *F. hepatica*

We performed scRNA-seq on cells from anterior and posterior parts of several adult individuals. First, we developed a protocol to dissociate anterior and posterior parts of the worms into single cells using a combination of mechanical and enzymatic treatment (Fig 2 A). Thereafter, viable cells were enriched by fluorescence-activated cell sorting (FACS) using calcein AM viability dye to select viable cells. By using the commercially available 10X Genomics Chromium platform, we analyzed a total of 10 samples, from either anterior (4) or posterior parts (4) of worms, or whole worms (2) for comparison. Using a combination of the 10X cellranger workflow and the R package Seurat^28^, we analyzed sequencing data of a final number of 19,581 cells. Hereby we detected a median gene number per cell of 2,644 and on average 14,087 UMIs per cell after quality filtering (suppl. table S1). Based on the total of 11,217 protein coding genes in the genome of *F. hepatica*, we detect on average 23% of the total genome as transcripts per cell. This is a touch higher compared to scRNAseq data obtained for adult *S. mansoni^19^*, where a median gene number of 1,600 was detected per cell, which is 16% of the total gene count of 9,896 protein coding genes in the genome of *S. mansoni^29^*.

**Fig 2.**
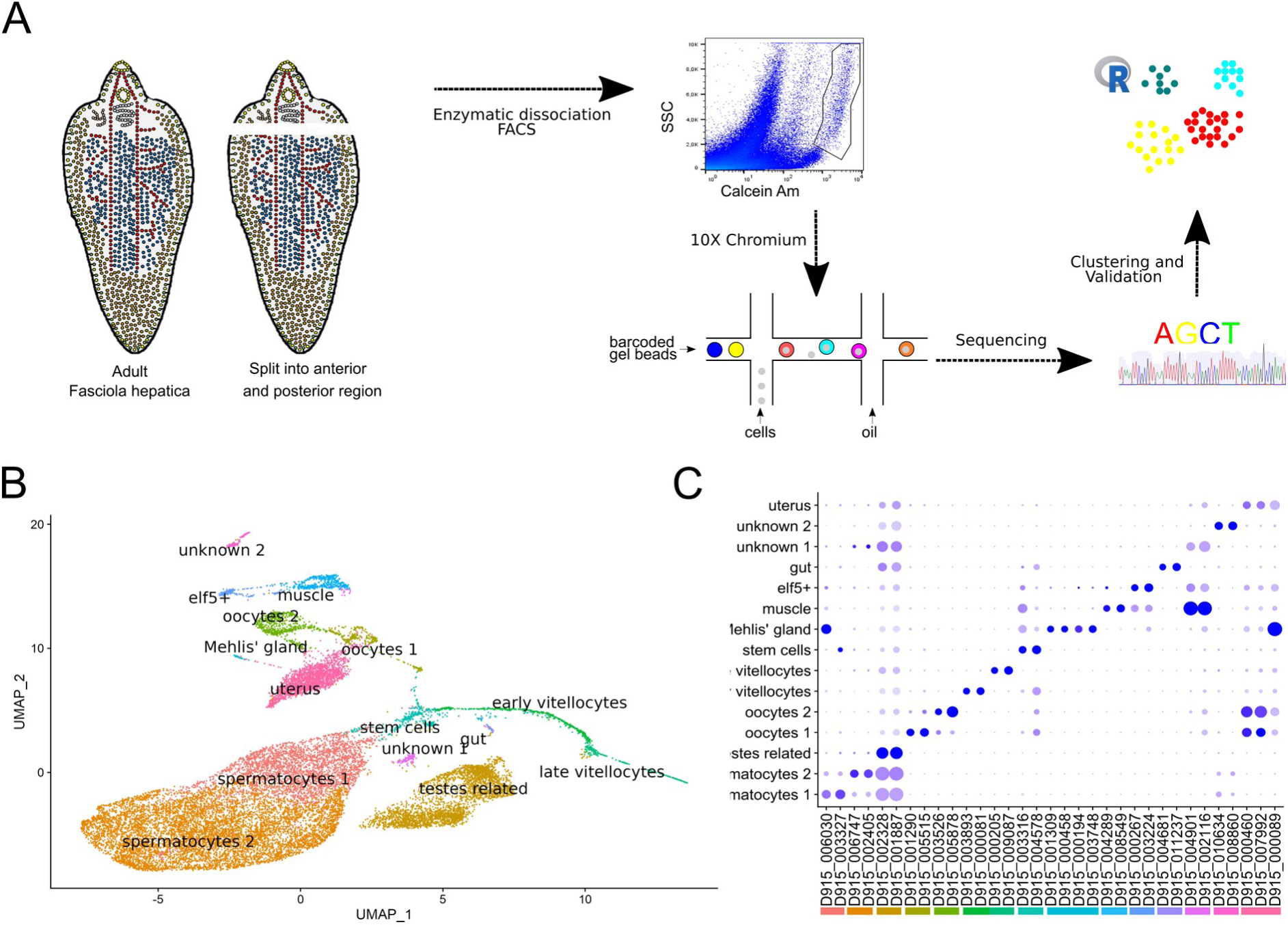
scRNA-seq allowed the classification of 15 cell clusters in adult *F. hepatica*. **A** Schematic workflow outlining the major steps for data generation: adult liver flukes were first split into an anterior and posterior part before dissociating into single cells by mechanical and enzymatic processing. Next, cells were sorted based on the viability dye calcein. Cells were barcoded following the 10X Chromium protocol. Libraries were sequenced and clustering was carried out to identify transcriptionally distinct cells. **B** Uniform Manifold Approximation and Projection (UMAP) of 19,581 cells. Clusters are colored and labels added. **C** Profiles of gene expression over all clusters illustrated as dotplot. Shown is the expression of at least two marker genes for each cluster, and the cluster color is indicated below each marker pair. Level of expression is indicated by color from blue (high expression) to lavender (low expression). The percentage with which the cells of a cluster express the given gene is represented by the size of the circles.

Using Seurat, we identified 15 clusters (Fig 2 B), for which distinct marker genes could be derived (Fig 2 C, suppl. table S2). The clusters were annotated with the help of published cell marker genes that are conserved across taxa, as well as by comparison to cellular markers for the closely related blood fluke *S. mansoni*. We identified clusters resembling cells of the following tissues (number of clusters in brackets): muscle (2), gut (1), testes (3), stem cells (1), ovary (2), vitellarium (2), eggs/uterus (1), Mehlis gland (1), and two clusters for which we could not find an annotation. These cell types, as hypothesized earlier, were not equally distributed between the anterior and posterior samples (suppl. Fig S1). Because gut, testes and vitellarium represent the largest tissues in *F. hepatica* and cover most of the posterior body part, the whole-worm samples and posterior samples strongly resembled each other. Specific cell types are missing in the data set, like parenchymal or neuronal cells, which could be explained by several factors. First, rare cell types might, despite our enrichment strategy, still be overrun by overabundant cell types and not be captured during scRNA-seq. Another reason may be linked to the chosen tissue dissociation protocol, which made use of enzymatic digestion followed by flow sorting, an established procedure for other flatworms^18,19^. One cannot fully exclude damage of more sensitive cells and/or a loss of syncytial cells, such as those of the tegument. Single-nuclei sequencing may be an elegant solution to this problem, but has so far never been applied to flatworms.

### Stem cells show distinct transcriptional profiles of proliferating and dormant states

Being a multicellular organism, the growth and development of *F. hepatica* depends on stem cells that give rise to progenitor cells of reproductive and somatic tissues^30^. The remarkable output of thousands of eggs per day^8^ necessitates a massive proliferative activity of germline stem cells and differentiation of germ cells. Understanding what drives that remarkable fecundity of the parasite involves understanding the gene expression controlling germline stem cells, i.e. spermatogonia and oogonia. First, the stem-cell cluster was identified based on the expression of known marker genes (Fig 3 A), like three nanos isoforms (D915_007877, D915_002111, D915_002112) previously described by Robb et al 2022^16^. These RNA-binding proteins are known regulators of neoblast function in different organisms, including flatworms^31–33^. Other marker genes for this cluster were histone 2b *(h2b*) (D915_007751) and several genes encoding MCM complex proteins. On closer inspection, we found that these marker genes showed two different types of expression patterns. While *nanos* expression was tightly restricted to the stem-cell cluster, other gene transcripts (e.g. *h2b*) were also detected in potential progenitor clusters of the vitelline, oogenesis and spermatogenesis lineages (Fig 3 D). Transcription of *h2b* was previously used to label actively proliferating cells in schistosomes and planarians^32,34^. To confirm that presence of *h2b* expression is a suitable marker for active cell proliferation in *Fasciola*, we labeled proliferating cells with the thymidine analogue 5-ethynyl-2’-deoxyuridine (EdU) and stained *h2b* transcripts with FISH. We found an overlap between *h2b* positive cells and EdU positive cells, with around 70% of EdU positive cells being also positive for *h2b* transcripts (Fig 3 C). The *h2b/*EdU double-positive cells were located in the periphery of the testicular lobes and ovary (Fig 3C), which validates prior description of stem cells in that location based on histological analysis^27^. Furthermore, *h2b* positive cells were located close to the tegument and gut tissue, confirming the presence of somatic stem cells in adult flukes, as described in previous studies for juvenile worms^30^. We also detected strong staining for *h2b* transcripts in the center of the ovary, containing mature oocytes. This can be explained by the fact that unlike stem cells, in which histone transcription is coupled to the cell cycle, both processes are decoupled during oogenesis in preparation of embryogenesis^35^.

**Fig 3.**
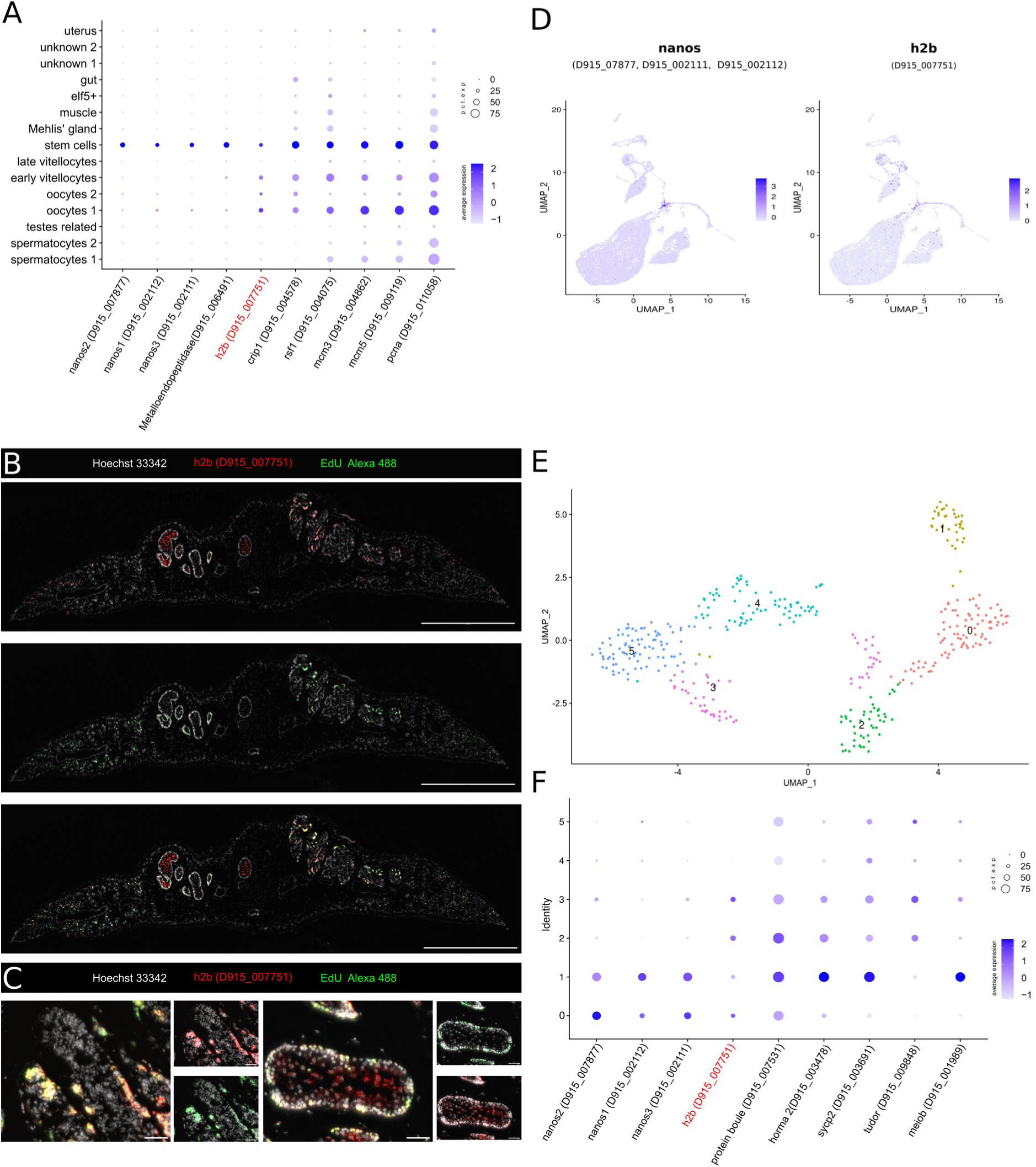
Stem cells show transcriptionally distinct cell states. **A** Dotplot showing the expression of stem-cell marker genes. ISH-validated genes are marked in red. Level of expression is indicated by color from blue (high expression) to lavender (low expression). The percentage with which cells of a cluster express the given gene is represented by the size of the circles. **B** Transversal section stained for *h2b* transcripts by FISH (red) and proliferating cells with EdU (green). Scale bar: 1000 µm. **C** Close-up of testis (left) and ovary (right) from B. Scale bar: 50 µm **D** UMAP plot showing the expression of *nanos* isoforms and *h2b*, **E** UMAP of subclustered cells colored by cluster. The clusters were numbered 0 to 6. **F** Dotplot showing the expression of *h2b*, *nanos* isoforms and marker genes for cell differentiation for the subclusters from E. ISH-validated genes are marked in red. Level of expression is indicated by color from blue (high expression) to lavender (low expression). The percentage with which cells of a cluster express the given gene is represented by the size of the circles.

To obtain a clearer picture on the differentiation dynamics of stem cells and their progeny cluster, we reanalyzed all cells within the stem-cell cluster and obtained six subclusters (Fig 3 E). Marker genes for several subclusters were enriched for genes involved in meiotic division (suppl. table S3). This included genes like *sycp2* coding for the synaptonemal complex protein 2, which was shown in immunolocalization experiments to be expressed in early progenitor cells of the reproductive organs of *F. hepatica^36^*, or the DAZ family member *boule*, which was shown in planarians and schistosomes to be important for germ-cell differentiation^20,37^. The six subclusters could be allocated to different cell differentiation states (Fig 3 E): prominent *nanos* and *h2b* expression with negligible expression of differentiation markers likely reflects actively proliferating stem cells (cluster 0); this gives rise to clusters that highly express both, *nanos* and differentiation markers (cluster 1), or mainly express differentiation markers (clusters 2 and 3). These clusters may cover stem-cell progeny on the way to germ-cell differentiation. While these previous clusters all express *h2b* to a certain extent, the two final, probably most differentiated clusters only express some differentiation markers (clusters 4 and 5). We thereby captured separate stages of differentiation in germline cells, which can aid in unraveling dynamics of germ-cell differentiation.

### Gene signatures of different cell states during germ-cell development

*F. hepatica* is a hermaphroditic flatworm, harboring both female and male reproductive tissues. We captured clusters of both tissues, which included cells related to more undifferentiated as well as more mature cells with distinct marker profiles (Fig 4 A). The testes are represented by three clusters (Fig 4 A). Two of the three clusters were successfully annotated based on their expression of the orthologs of described marker genes of the male germline in schistosomes as well as markers predicted from the subclustering of the proliferating cells. Cells in the spermatocytes 1 cluster expressed *boule* (D915_007531) and the transcription factor *one cut 1* (D915_002483), both of which are described to promote male germ-cell differentiation^20,37^. Additional marker genes also hinted at cell proliferation and the initiation of differentiation as main processes in this cluster, like genes coding for histones or RNA helicases. Furthermore, we additionally found a strong expression of a gene annotated as the meiosis specific with OB-fold (*meiob*) gene, which is known to play a role in meiotic recombination in humans^38,39^, within the spermatocyte 1 cluster. Expression of *meiob* within the testes was confirmed by ISH (Fig 4 B). In contrast to this, the spermatocyte cluster 2 was enriched for genes encoding structural components like several tubulins and tektins known to be part of the sperm flagellum (Fig 4 B). One of the tektin genes was also shown to be strongly expressed in the testes of the worm by ISH (Fig 4 C). Further support for the functions associated with each cluster was obtained by GO term analysis (suppl. table S4). The spermatocytes 1 cluster was enriched for GO terms like RNA binding or nucleotide binding as well as several metabolic processes, further underlining a more proliferative cell state. In contrast, the spermatocytes 2 cluster was enriched for terms involved in cytoskeletal organization, cilium assembly and axoneme assembly (suppl. table S4), reflecting a differentiated cell state^27^.

**Fig 4.**
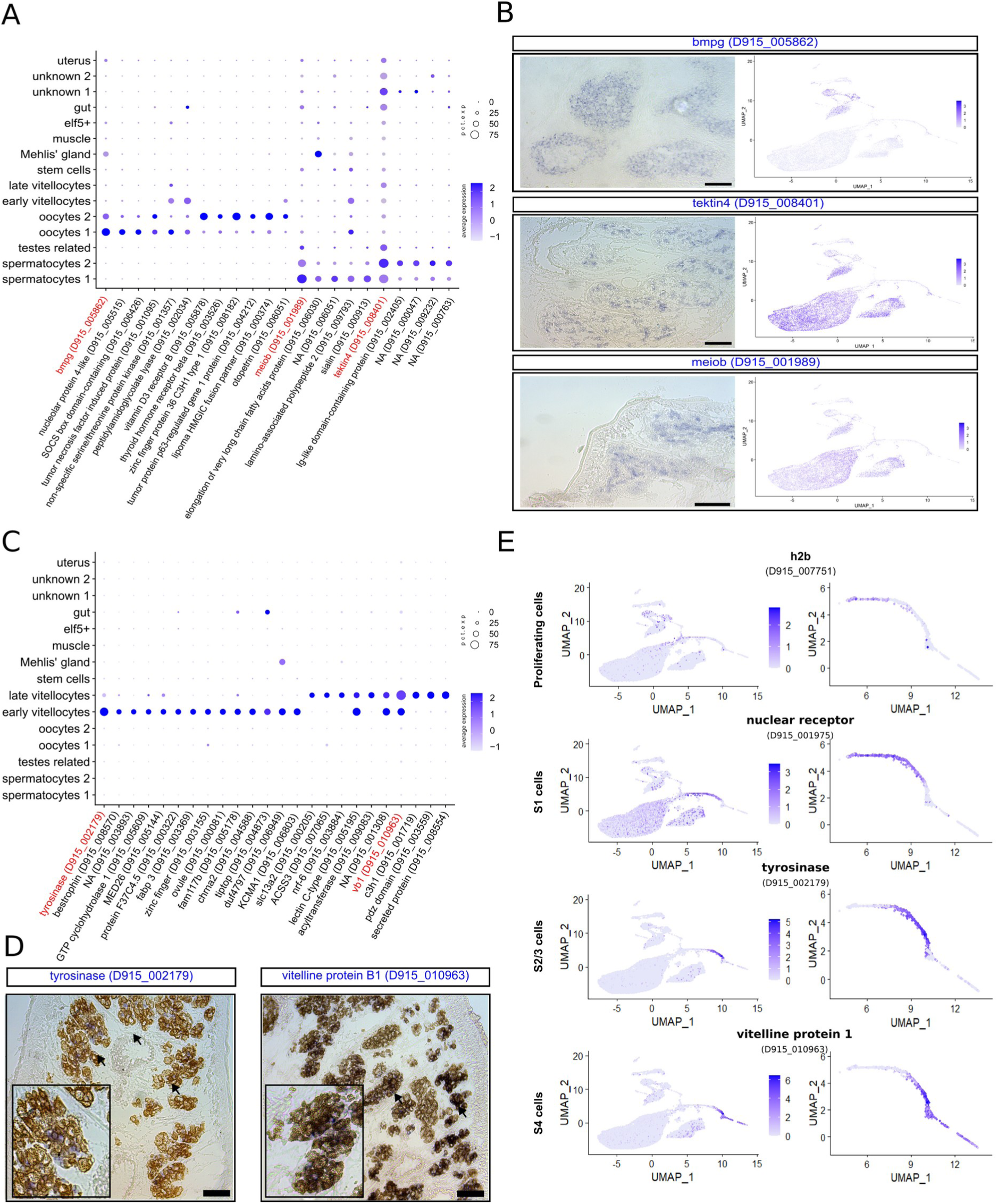
Conserved marker genes in cells of reproductive tissues including a vitelline cell lineage. **A** Dotplot showing the expression of germ cell marker genes. ISH-validated genes are marked in red. Level of expression is indicated by color from blue (high expression) to lavender (low expression). The percentage with which cells of a cluster express the given gene is represented by the size of the circles. **B** ISH stainings (top row) for transcripts of selected male and female germline markers *bmpg*, *tektin4* and *meiob* (in blue), and corresponding UMAP plots of gene expression. Scale bar: 100 µm. **C** Dotplot showing the expression of vitelline cell marker genes. ISH-validated genes are marked in red. Level of expression is indicated by color from blue (high expression) to lavender (low expression). The percentage with which cells of a cluster express the given gene is represented by the size of the circles. **C** ISH staining for the vitelline cell markers *tyrosinase* and *VB1* transcripts (blue), arrows indicate positive staining. Scale bar: 100 µm. **D** UMAP plots showing the expression of conserved vitelline-cell marker genes shared with vitelline-cell markers of *Schistosoma mansoni^42,43^*. An overview of all clusters and close-up of the stem cell, early and late vitellocytes clusters is shown.

The ovary of *F. hepatica*, as the testes, is a branched organ consisting of three major cell types, the proliferating oogonia and the differentiated oocytes, which are further classified in oocytes 1 and 2, based on their differentiation state^36^. We identified two clusters of the ovarian cells based on their expression of the bone marrow proteoglycan (*bmpg*), a gene characteristic for oocytes found in schistosomes^19^. Transcripts of the *F. hepatica bmpg* localized to the more mature oogonia in the center of the ovary in ISH experiments (Fig 4 B). As outlined before, late and early oocyte clusters differed in their expression of synaptonemal complex proteins^36^. As for the spermatocyte 1 cluster, GO terms related to metabolic processes, DNA binding or DNA replication were enriched in the early oocyte cluster. Interestingly, the terms enriched within the late oocyte cluster were related to processes like signaling, phosphorylation, organization and nuclear receptor activity (suppl. table S4), emphasizing a complex control of signaling processes in these cells.

### Differentiation markers of vitellocytes are conserved across species

The vitellarium is a unique organ of flatworms, essential to produce vitellocytes that enter ectolecithal eggs together with one oocyte. Vitellocytes (vitelline cells) are important for egg shell formation and contain nutrients for the later development of the embryo. In *Fasciola*, like in schistosomes, the immature vitelline cells heavily proliferate and develop in four different stages^40,41^. While vitellocytes and their gene expression have seen interest in schistosome research^19,42–44^, no such data was available for *F. hepatica* and the different cell states have been mainly categorized by their morphology^40^. We discriminated two main clusters based on their differential gene expression, which we named early and late vitellocytes. Identification of orthologs for typical genes characterizing S1 to S4 vitelline cell stages in *S. mansoni^44^* allowed us to display the full vitelline cell lineage in *F. hepatica*. The ortholog for the nuclear receptor vitellogenic factor 1 (D915_001975) was expressed in the early vitellocytes cluster (Fig 4 E), partially coexpressing the proliferative marker *h2b*. Expression of tyrosinase (D915_002179) characterized an intermediate state (S2/S3) between the early and late vitellocyte clusters (Fig 4 E), which also agrees with earlier observations in schistosomes, where maturing cells express tyrosinases but expression is absent in mature S4 cells. The expression of this tyrosinase was additionally validated in the vitelline cells by ISH (Fig 4 D). Finally, the expression of egg-shell genes, such as *vb1* (D915_010963)^45^, was found to be specific for mature S4 vitellocytes. As expected, GO terms enriched in the early vitellocyte cell cluster were related to proliferation and transcriptional activity, like nucleotide binding or RNA processing, while terms within the late cells covered terms like iron or vitamin binding (suppl. table S4). Overall, the existence of shared molecular vitelline-cell markers for blood and liver flukes suggest that the biological mechanisms guiding vitelline-cell function and maturation are conserved. Also major differentiation markers of vitelline cells appear conserved across these two major trematode species. This degree of conservation is anything but self-evident, given the highly differential sexual biology of both pathogens – with liver flukes being hermaphrodites and schistosomes dioecious. Insights in the reproductive biology of flatworms may allow the development of strategies to limit transmission of the parasites, consequently, the vitellarium has been discussed as a valuable target^46^.

### Genes involved in lipid metabolism characterize gastrodermal cells of liver flukes

As for most trematodes, the intestine of liver flukes is bifurcated with numerous branches stretching throughout the parasite’s body. The gastrodermal cells are known to express and secrete a high number of digestive enzymes, primarily cathepsins^47–49^. Accordingly, we classified cells expressing known intestinal cathepsins of *F. hepatica^12^*(suppl. table S5) as gastrodermal cells (Fig 5 A). We validated this by ISH of cathepsin L1 (Fig 5 D). As to be expected, a high number of characteristic genes for this cluster associated with GO terms like proteolysis, cysteine-type peptidase activity or lipid binding (Fig 5 B). Related to lipid metabolism, a gene coding for a phospholipase B-like protein (D915_003832) caught our attention, which showed specific transcript staining in the branches of the gut (Fig 5 B). It can be speculated that this gut-specific protein may be part of lipid metabolism in gastrodermal cells of the adult worm. As liver flukes have a highly reduced lipid metabolism^50^, processing of endogenous or host-derived lipids in gastrodermal cells warrants future investigation and could be interesting as anthelminthic target.

**Fig 5.**
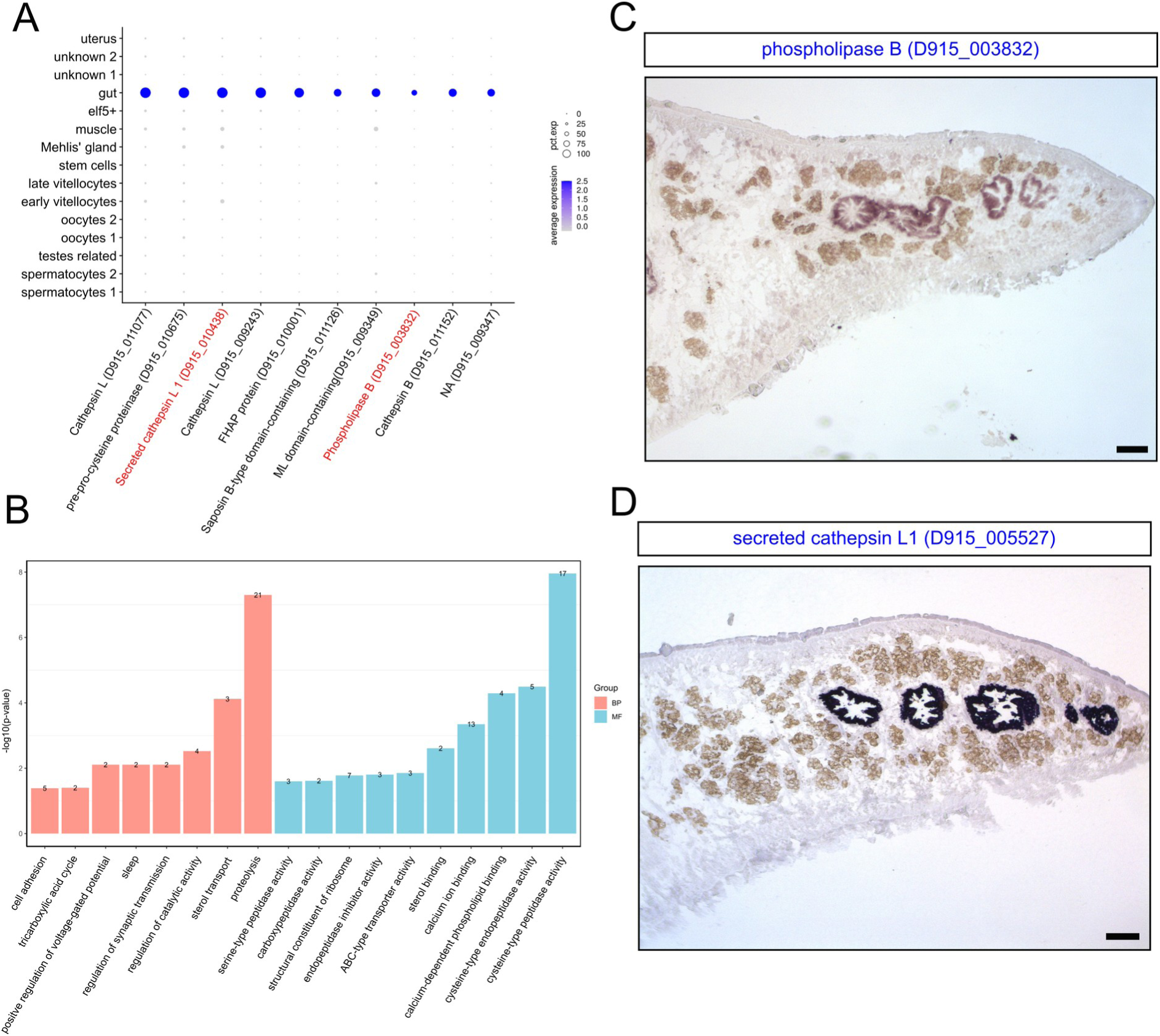
Gut cells of *F. hepatica* express genes involved in lipid metabolism. **A** Dotplot showing the expression of gut cell marker genes. ISH-validated genes are marked in red. Level of expression is indicated by color from blue (high expression) to lavender (low expression). The percentage with which cells of a cluster express the given gene is represented by the sizes of the circles. **B** Gene ontology analysis of marker genes (top 75% per cluster) revealed characteristic biological processes (BP) and molecular functions (MF). The number of enriched genes is noted at the end of each bar. **C** ISH staining for *phospholipase B* transcripts and **D** for *cathepsin L1* transcripts in the gastrodermis (blue to dark blue color). Scale bar: 100 µm.

### Specialized muscle cells express several kinase genes

The musculature in trematodes is important for several functions essential for the fluke’s survival, like attachment to the bile duct to keep in place, or the feeding activity. The muscular system is composed of body musculature controlling the worm’s movement, the sucker musculature for attachment, as well as the muscles lining the reproductive and digestive organs^51^. All muscles are of the invertebrate smooth type with the cell body connected to the muscle fiber via cytoplasmic connections^46,52^. The liver fluke musculature was represented by two clusters in our data, which were termed muscle and elf5+. The muscle cluster was identified by the expression of myosin and collagen as marker genes (Fig 6 B) as well as conserved muscle markers from other species^53^. The expression of collagen in muscle cells is in agreement with studies in planarians and schistosomes^19,54^, which described muscle cells as a main source of extracellular matrix. We further validated the expression of collagen and myosin by ISH. As a reference, we combined FISH with immunofluorescence, utilising an antibody against muscle fiber protein frequently used in planarian research^55^. Cells expressing collagen and myosin were widley distributed and located in close proximity to the stained muscle fibers in both the subtegumental muscle layer and throughout the body (suppl Fig S3). This pattern clearly fits the nature of invertebrate muscle fibers described above. Next to these structural molecules, we found expression of a gene coding for the 5-HT receptor 1 (D915_001848), (Fig 6 A), which is thought to be involved in the serotonin-dependent activation of muscle fibers in flatworms^56,57^, as well as other central regulators of cell signaling. Among these were protein kinase C, G-protein coupled receptors and phosholipase C, for which signaling in response to FMRFamides was previously suggested for *Fasciola* muscle fibers^58^. Unexpectedly, expression of the cystatin like stefin-2 (D915_009861) was high within the muscle cluster. Previous work on *F. hepatica* showed the localization of stefins within gastrodermal cells, the tegumental area as well as the reproductive organs^59^. Our data suggest that the transcripts for stefin-1 (D915_009335) were indeed present in cells of the reproductive organs as well as in the gut, while stefin-2 (D915_009861) and stefin-3 (D915_001085) were also present in the muscle cluster, which is the first time the expression of those protease inhibitors is described in this cell type (suppl. Fig S2).

**Fig 6.**
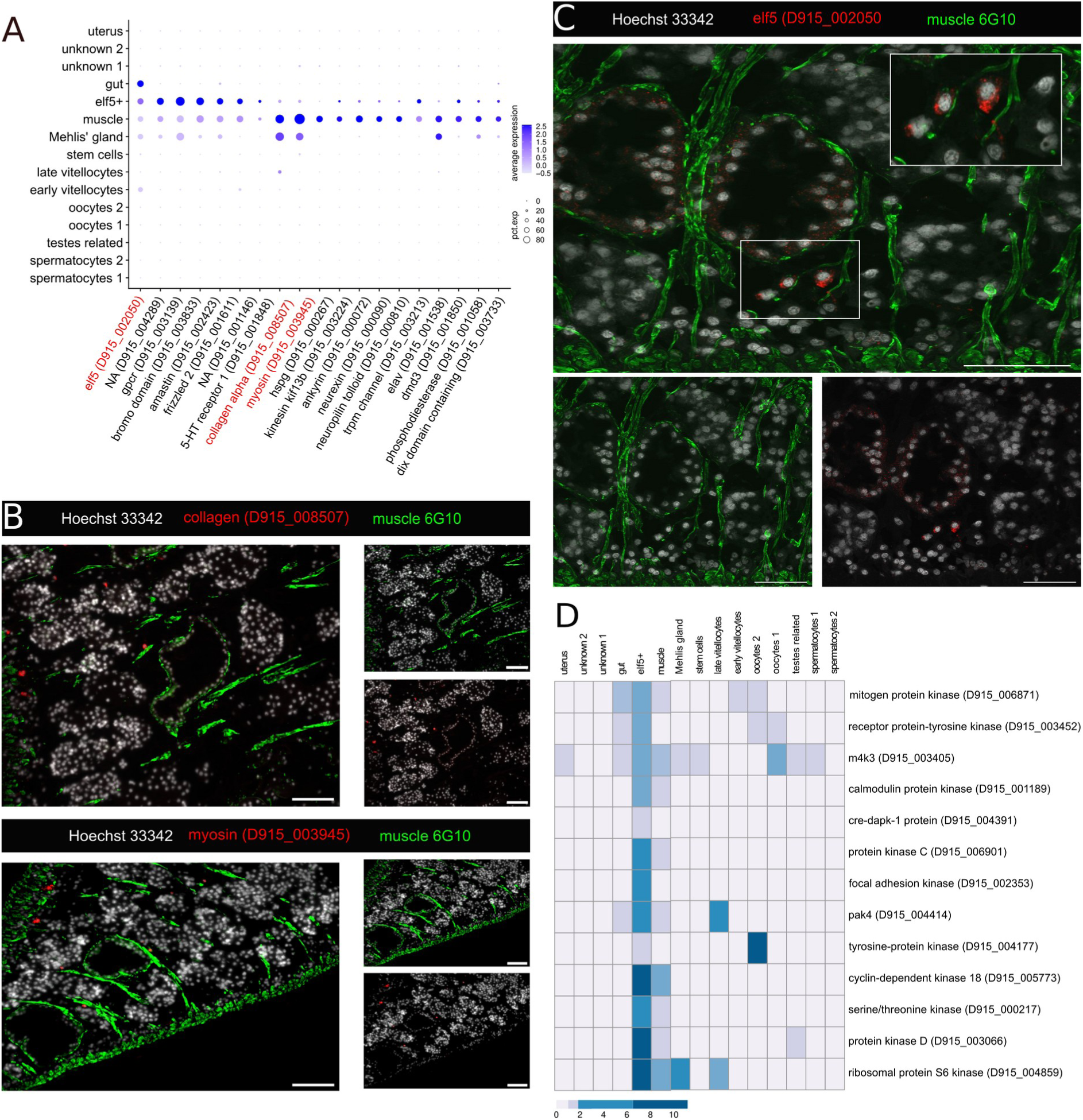
Specialized muscle cells express several protein kinase genes. **A** Dotplot showing the expression of marker genes for the muscle and elf5+ muscle cell clusters. ISH-validated genes are marked in red. Level of expression is indicated by color from blue (high expression) to lavender (low expression). The percentage with which cells of a cluster express the given gene is represented by the size of the circles. **B** Detailed view of FISH for collagen (top) and myosin (bottom) combined with immunolocalization of muscle fiber proteins. Scale bar: 100 µm. **C** FISH for elf5 combined with immunolocalization of muscle fiber proteins. Scale bar: 100 µm **D** Average expression of protein kinase marker genes per cluster displayed as a heatmap. Note the high and enriched expression of several protein kinases in the elf5+ muscle cluster. Expression values were centered and scaled for each row (each gene) individually.

An additional cluster had muscle-like features based on marker overlap with the muscle cluster, though it also expressed distinct marker genes (Fig 6 A). We termed this cluster elf5+ based on the highly characteristic expression of the ets transcription factor elf5 (D915_002050), which was shown to regulate extracellular matrix composition in planarians^60^. In order to locate these elf5+ cells within *F. hepatica*, we performed FISH. Cells positive for *elf5* transcripts located in close contact with muscle fibers (Fig 6 C), which further underlined their annotation as muscle cells. While the muscle cluster appeared to be more involved in metabolic processes, translation and transport, the elf5+ cluster was enriched in GO terms related to cell communication and signal transduction. Related to these terms are for example the neuropilin and tolloid protein (D915_000810), which has been found in neurons as well as a type of smooth muscle cells in humans^61^ and functions in the neuromuscular junction in *Drosophila melanogaster^62^*, or two genes enocoding for ankyrin 2 (D915_000071, D915_002954), which contain an ankyrin repeat region. The family of ankyrins is involved in the attachment of membrane proteins to the cytoskeleton, and especially type 2 ankyrins have been shown to function in muscle cells^63,64^.

To characterize this cell type in more detail, we focused on protein kinases, which are central regulators of a plethora of cellular process^23^. A notable number of 16 PKs was found among the marker genes (Fig 6 E and suppl. Table S2). These included two proposed protein kinase C (PKC) genes, D915_006901 and D915_003066, though while annotated as PKC, D915_003066 shows closer sequence similarity to protein kinase D. PKC signaling was previously discussed as mediator of body contraction in schistosomes and liver flukes based on studies with whole worms or body strips of worms treated with PKC activators or inhibitors^58^. Our expression data now proof PKC expression in muscle cells and thereby suggest an involvement of PKC signaling in muscle function^65,66^. Other highly characteristic kinases were the focal adhesion kinase (D915_002353) and the p21-activated kinase 4 (PAK4, D915_004414). These kinases are typically involved in signaling related to extracellular matrix binding and focal adhesions^67,68^. Another interesting feature of the elf5+ cluster is the high expression of the TRPM_PZQ_ channel (D915_003213) (Fig. 6A), hinting at the still unknown role of this channel in parasitic flatworms. We speculate on a role in cell adhesion, as TRPM channels have shown to localize in focal adhesions in humans^69^. Ligands targeting the schistosomal TRPM_PZQ_ (praziquantel) or the *Fasciola* TRPM_PZQ_ (BZQ) caused tegumental damage^70,71^, which might be related to the destabilization of the attachment of cells to the extracellular matrix, and to the basement membrane in particular.

Taken together, scRNA-seq analysis revealed a previously unrecognized muscle cell type that may have a more specialized location and role, thereby being distinct from the main subtegumental musculature.

### An inhibitor of the p21-activated PAK4 kinase reduces fluke vitality

Protein kinases (PKs) are well druggable targets, also in various helminths^72^. The important role of PKs in the formation and progression of cancer has led to the development of small-molecule inhibitors, which have potential for drug repurposing against parasites^21^. The enrichment for various kinases within the elf5+ cluster as well as the expression of the pan-flatworm target TRPM_PZQ_ led us to the hypothesis that the cells within the elf5+ cluster might be valuable for the identification of novel drug targets with vital functions for the parasite.

We selected PAK4 as a candidate kinase. PAK4 represents a still unexplored kinase in trematodes, but has shown promise in the treatment of various forms of cancer^73^. PAK4 is a member of the p21-activated kinase (PAK) family, which is characterized by a p21-binding domain (PDB), also called Cdc42/Rac interactive binding (CRIB) motif^74^. In humans, this kinase family is involved in several developmental processes, functions in the immune system and the nervous system^75^. In the fruit fly *D. melanogaster*, these proteins play roles in the development and control of the visual system^76^, while for *Caenorhabditis elegans,* they are important for axon guidance, gonad development and mechanotransduction^77–79^. To date, PAK kinases have not been addressed in parasitic helminths. First, we confirmed the presence of the essential protein domains within the FhPAK4 amino acid sequence by SMART analysis (suppl. Fig S3). FhPAK4 shared overall 44.6% sequence identity with human PAK4, while their kinase domains had 64.3% sequence identity. Most organisms have multiple PAK family members, which belong to two groups. Group 1 and group 2 differ in their structure as well as their function^74,80^. Within the genome of *F. hepatica*, we detected three more PAK kinases next to FhPAK4, based on the presence of the characteristic PDB domain. By comparing *F. hepatica* PAK sequences with sequences from model organisms, FhPAK4 could be identified as a group 2 family member, while the other PAKs allocated to group 1 (Fig 7 A). Based on these results, we termed D915_001478 and D915_004654 as FhPAK1 and FhPAK2, respectively. The last ortholog, D915_006992, clustered within a previously described group^77,81^ together with the more divergent PAK members *D. melanogaster* PAK3 and *C. elegans* MAX-2, which is why we named this kinase FhPAK3. While *fhpak1* and *fhpak2* were transcribed in several different cell types, *fhpak3* was expressed abundantly in mature oocytes and *fhpak4* expression was confined to the elf5+ cluster, the gastrodermis and vitellocytes (suppl. Fig S4). FISH experiments localized *fhpak4* transcripts in gastrodermal cells, in oocytes, and in large, scattered muscle cells positioned clearly distant from the typical subtegumental body musculature (Fig 7 B) as seen for *elf5* transcripts. The presence of *fhpak4* transcripts within the nucleus of oocytes might hint at a role for FhPAK4 in early embryo development. In line with this hypothesis is the high abundance of transcripts within the eggs of *F. hepatica* that we found in available bulk RNA-seq data (Cwiklinski 2015). Maternal transcripts of *pak4* were also detected in the developing embryo of zebrafish, where PAK4 is essentiell for myelopoisis^82^.

**Fig 7.**
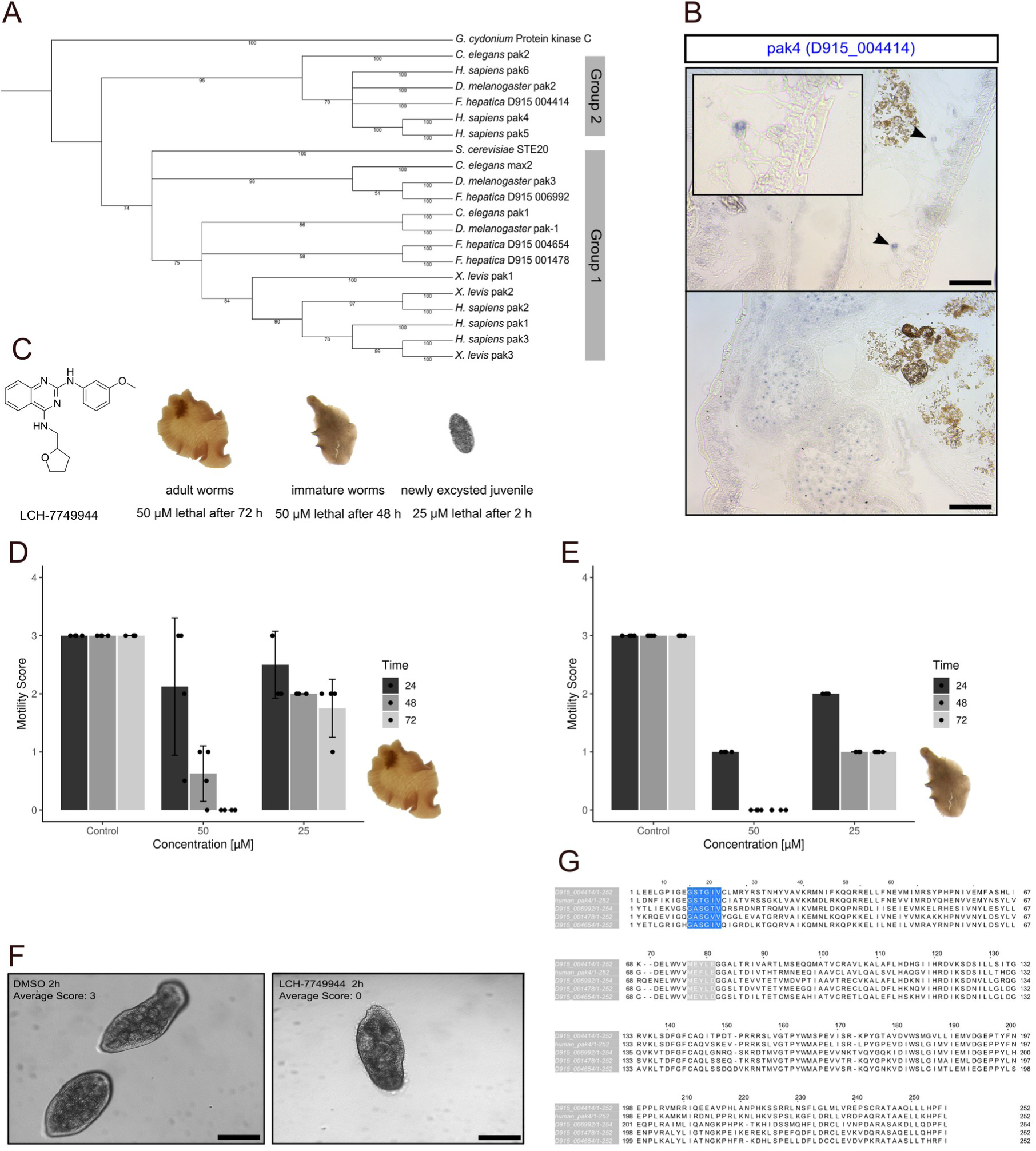
An inhibitor of the p21-activated PAK4 kinase reduces parasite vitality. **A** Phylogenetic tree of PAK4 orthologs of *F. hepatica* and other species (accession numbers see suppl. table S6). p21 families are indicated, protein kinase C from *Geodia cydonium* served as outgroup. **B** ISH staining for Fh*pak4* transcripts. Scale bar: 100 µm. **C** Chemical structure of the PAK4 inhibitor LCH-7749944 with overview of lethal effects on different liver fluke stages. **D, E** Motility scores of adult worms (D) and immature worms (E) under varying LCH-7749944 concentrations over 72 h (4 replicates per condition). Error bar shows standard deviation. **F** NEJs after 2 h treatment with 50 µM of LCH-7749944 (right) compared to vehicle-treated NEJs (left). Representative images of 3 worms per condition are shown. Scale bar: 200 µm. **G** Alignment of human PAK4 amino acid sequence with *F. hepatica* PAK sequences. Binding sites of LCH-7749944 are colored: P-loop in blue and hinge region in gray.

Due to the numerous roles of the human PAK4 in various cancer types, small molecule-inhibitors have been developed^83,84^. We evaluated the efficacy of the ATP-competitive inhibitor LCH-7749944, which was shown to have an impact on proliferation, migration and invasion of gastric cancer cells^85^. The binding mechanism of LCH-774994 was predicted to involve interaction with residues within the hinge/p-loop regions of the human PAK4^86^. We found high sequence similarities between human and FhPAK4 for both regions, while the three other *Fasciola* PAK proteins diverged to a higher degree (Fig 7 G). This suggests that LCH-774994 may target specifically FhPAK4 in *F. hepatica*. To identify the potential of FhPAK4 as drug target, we tested this inhibitor against different disease-relevant parasite stages: newly excysted juveniles (NEJs) found in the host’s intestine, immature worms migrating and feeding through liver tissue, and the bile duct-residing adult worms. For all three stages, we observed a significant reduction of the motility during *in vitro* culture after LCH-774994 treatment (Fig 7 C). Adult worms showed moderate reduction in motility after incubation with 25 µM, but were severly affected with 50 µM after 48 h, ending in 100% lethality after 72 h (Fig 7 D). For immature worms, treatment led to a strong reduction of motility with 50 µM already after 24 hours, and all worms died after 48 h (Fig 7 E). Thus, immature worms appeared more susceptible compared to adults. NEJs responded even more sensitive, since all died within few hours of exposure to 25 µM, displayed impaired tissue integrity and prominent lesions on their tegument (Fig 7 F). Previous research highlighted the druggability and role of PAKs in human host cells, for example when it comes to host-cell invasion or pathogen-induced manipulation of the host‘s cytoskeleton by viruses, bacteria and parasites^87^. Here, we provide insight that PAK also of pathogen origin is essential for pathogen survival and appears attractive for therapeutic approaches. Thus, the example of FhPAK4 illustrates that the elf5+ muscle cells may be of high interest with respect to drug development.

## Conclusions

In this study, we present the first transcriptome for *F. hepatica* at single-cell resolution. The scRNA-seq dataset covers several cell types that are important for proliferation, as well as somatic cell types important for parasite vitality, like gastrodermal cells and cells of the musculature. The identification of molecular markers characteristic for each cell type delivered new information on cell lineages and cell-type specific functions. We described a previously unrecognized muscle cell type, characterized by expression of TRPM_PZQ_ and a multitude of protein kinases. By prioritizing the family of p21-activated kinases within this cell type, we highlighted the usefulness of this scRNA-seq dataset in the discovery of novel druggable targets. Thus, this dataset can serve as a resource for addressing basic and applied research questions, by providing valuable insights in the cellular biology of a multicellular pathogen.

## Limitations of the study

We report the transcriptional characterization of several cell types. Due to the abundance of cells of the male reproductive tract, this dataset misses some of somatic cell types, like cells of the parenchyma or the nervous system. Alternative ways to enrich for such cell types would add to this dataset.

## Acknowledgements

Financial support by the Deutsche Forschungsgemeinschaft (DFG) under grant HA 6963/2-1 and by the State of Hesse, LOEWE Center “DRUID”, is gratefully acknowledged. O.P. received a scholarship by the Justus Liebig University Giessen.

## Author contributions

Conceptualization, O.P., S.H.; Methodology, OP.; Investigation, S.G., O.P., S.A., JK..; Visualization, O.P.; Writing – Original Draft Preparation, O.P.; Writing – Review & Editing, all authors; Funding Acquisition, C.S., S.H; Supervision: SH.

## Supplemental figure titles and legends

**Fig S1.**
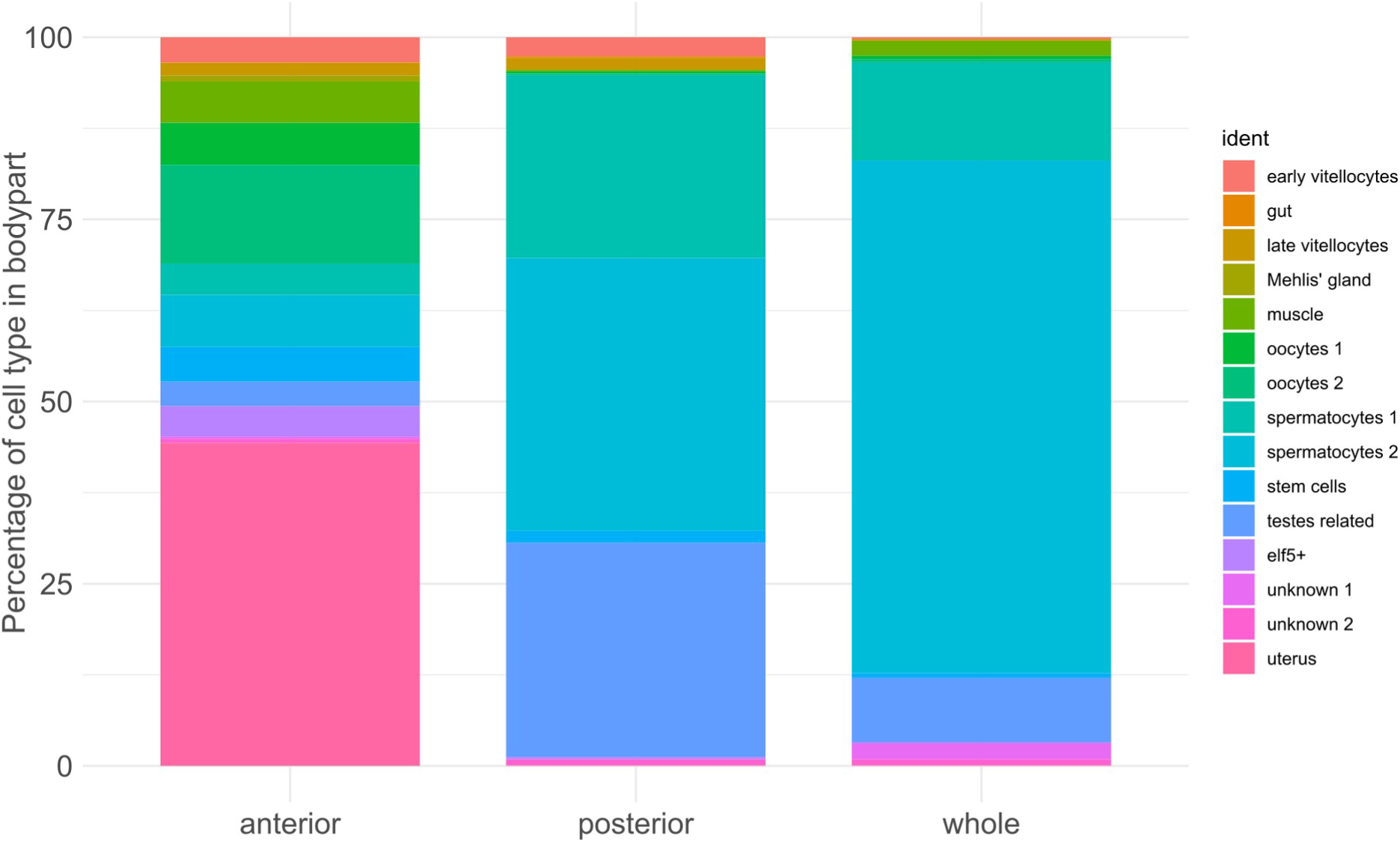
Cellular composition differs between samples. For each sample type - either anterior, posterior or whole worm sample - the number of cells within each of the 15 clusters was computed. Shown is the percentage that each cluster covers within the total cell number, per sample type.

**Fig S2.**
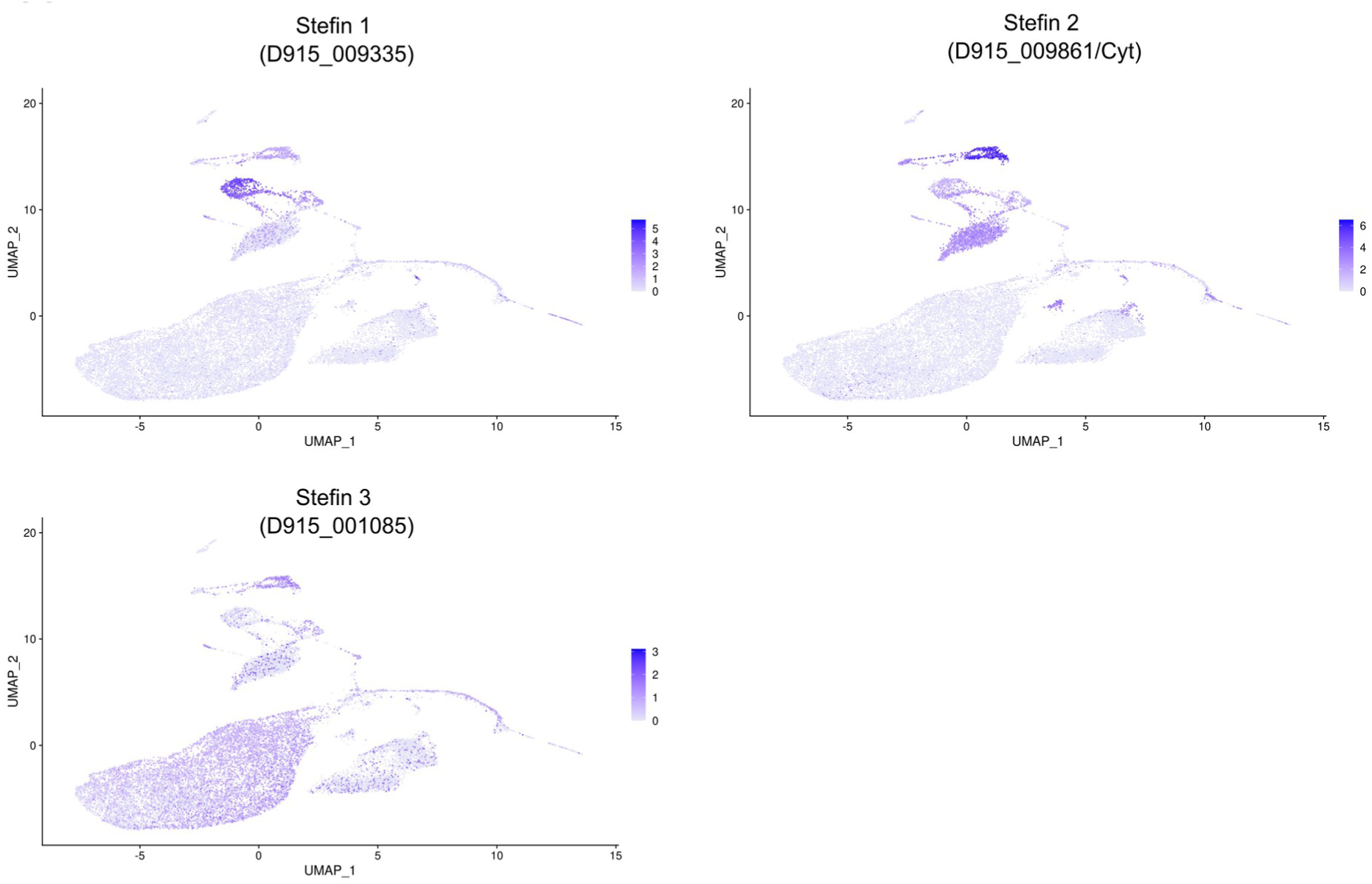
Different stefins show distinct expression patterns for different cell types. UMAP plots colored by gene expression showing the expression of three different stefins.

**Fig S3.**
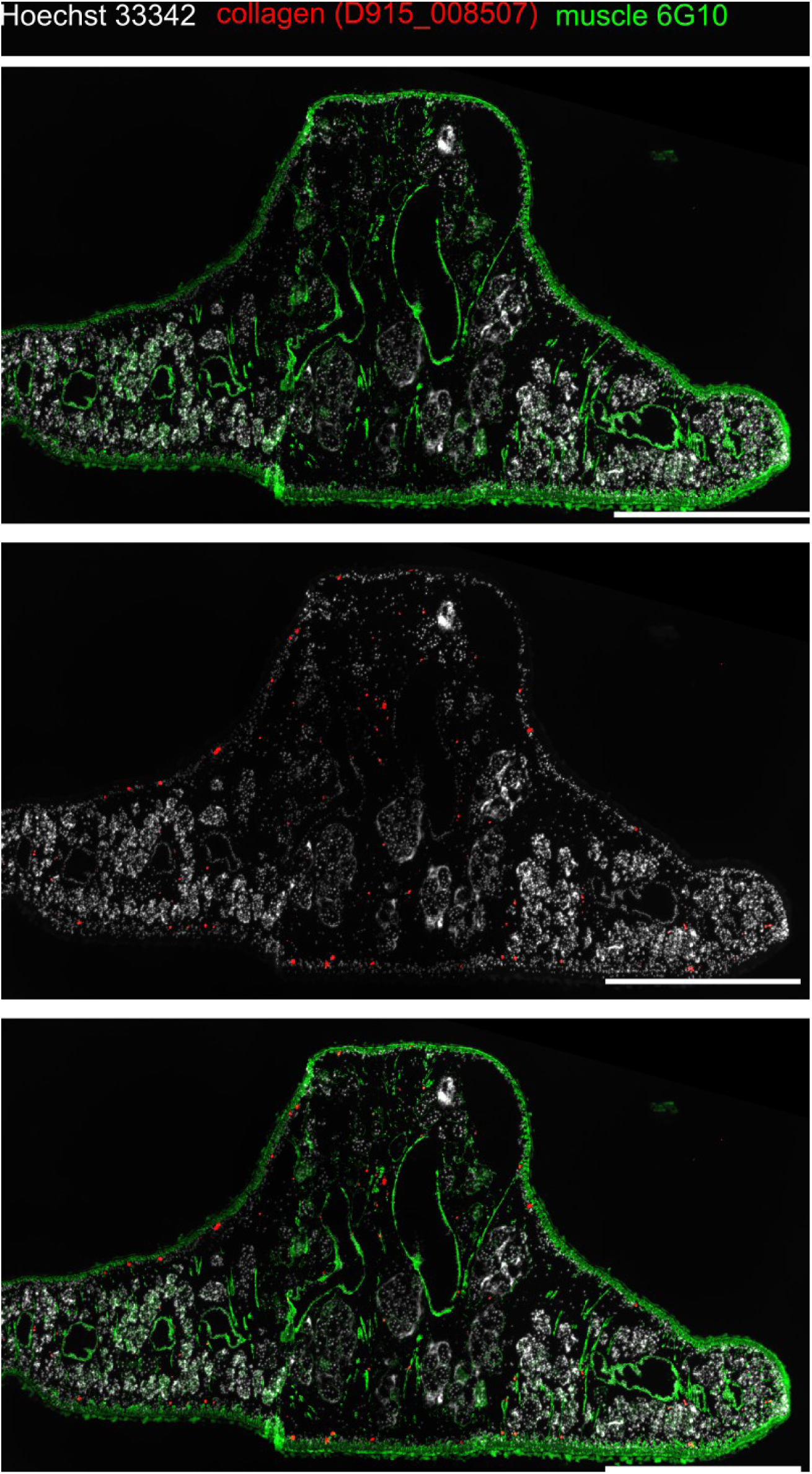
Muscle fibers and cells positive for collagen transcripts. Transversal section co-stained for collagen transcripts by FISH (red) and muscle fiber proteins (green) by immunolocalization. Scale bar: 1000 μm.s

**Fig S4.**
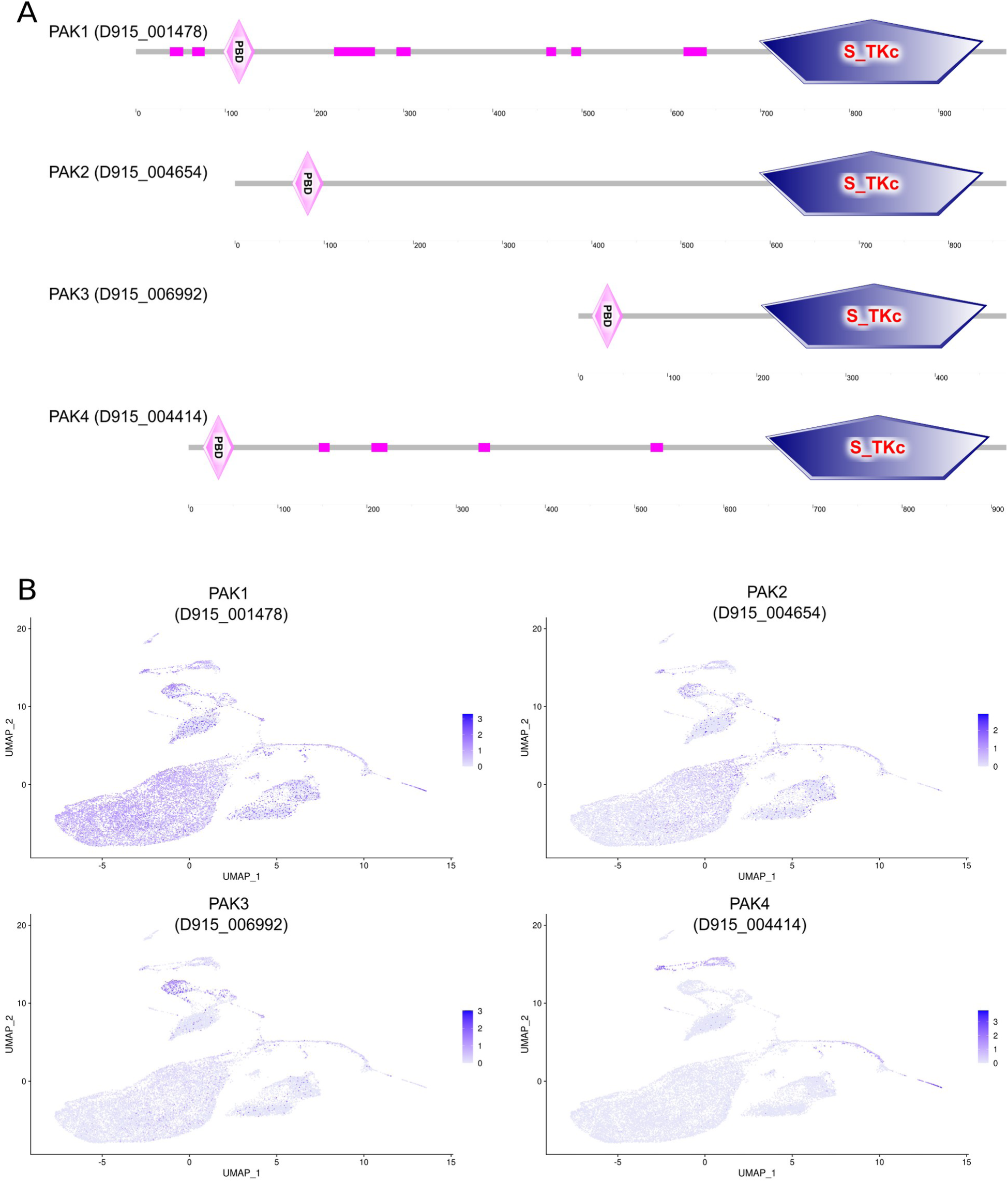
Domain structure and gene expression patterns of the four different PAK kinases. **A** SMART analysis confirmed the typical domain structure of *F. hepatica* PAK proteins, consisting of a serine/threonine kinase domain (S_TKc) and a p21-binding domain (PBD). **B** UMAP plots colored by gene expression showing the expression of various PAK genes.

## Supplemental video and Excel table titles and legends

**Table S1. List of samples used and related QC metrics after filtering**

**Table S2. List of marker genes per cluster**

The FindAllMarkers() function embedded in Seurat was used to identify markers for each of the clusters by “ROC” test

**Table S3. List of marker genes per stem cell subcluster**

The FindAllMarkers() function embedded in Seurat was used to identify positve markers for each of the clusters by “wilcox” test

**Table S4. GO terms enriched in clusters**

**Table S5. List of cathepsins used for definition of the gut cluster**

**Table S6 List of sequences used for phylogenetic analysis**

**Table S7 Primers used for cloning**

